# Extreme miniaturization in plasmid design: generation of the 903 bp cloning vector pICOt2

**DOI:** 10.1101/2023.11.29.569326

**Authors:** Jens Staal, Rudi Beyaert

## Abstract

Inspired by the size optimizations in the recently published pUdO series of plasmids, we set out to break our previous world record (pICOz, 1185 bp) in designing the smallest possible self-replicating high copy cloning vector. By replacing the zeocin resistance marker with trimethoprim resistance, deleting the parts upstream of -35 in the ampicilin resistance promoter and replacing the extended multiple cloning site with a single SpeI site, we managed to reduce the plasmid size to 903 bp, almost a 24% reduction in size compared to our previous accomplishments.

## Introduction

Following the minimalist philosophy expressed by the French poet Antoine de Saint Exupéry [1], [2] : *“It seems that perfection is attained not when there is nothing more to add, but when there is nothing more to remove”*, we have set out to generate the most minimal cloning vector possible. We believed that we had reached the absolute minimal possible vector in the generation of the 1185 bp pICOz vector [3]. However, the recently published pUdOs series of plasmids and another series of plasmids utilizing the tiny trimethoprim resistance genes [4], [5] showed that there are further opportunities for miniaturization. Inspired by this, we set out to investigate how much further we could miniaturize our previous pICOz cloning plasmid.

## Results and Discussion

### Generation of pICOt and pICOt2

We generated a 923 bp pICOt template using a synthetic design approach as described in the Materials and Methods section and compared this to the even smaller 903 bp pICOt2 generated by PCR from the pICOt template, where only the minimal part of the AmpR promoter remains (Fig. 1). For cloning, both pICOt and pICOt2 were amplified by PCR, digested and ligated prior to transformation in to *E. coli*.

**Fig. 1.**
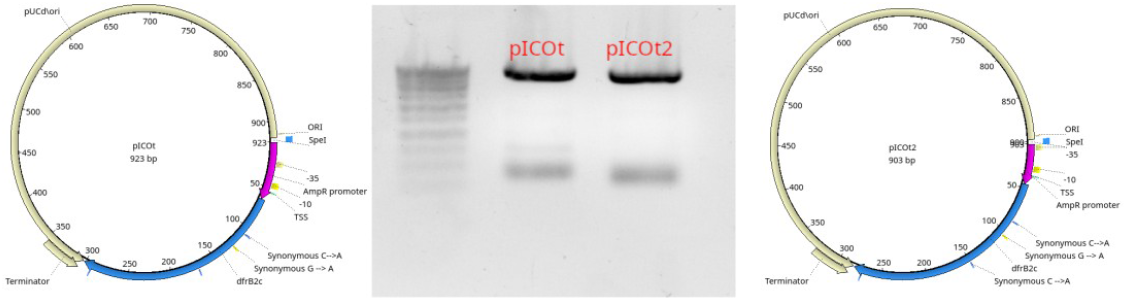
Design and PCR amplification of the complete minimal pICOt and pICOt2 plasmids (pink: Amp promoter; blue: trimethoprim resistance gene; yellow: minimal pUC ori). The PCR products of full-length pICOt and pICOt2 were run on a 1% Agarose/TAE gel and compared to the Eurogentec SmartLadder SF where the top band represent 1 kb.

The resulting plasmids were grown in normal LB or low-salt LB with trimethoprim selection. Selection with trimethoprim even at high concentrations (100 μg/ml) in normal LB was unable to completely block growth of mock-transfected MC1061 *E. coli* cells, whereas selection in low-salt LB was efficient at 50 μg/ml. The two plasmids pICOt and pICOt2 grew equally well with similar yields, indicating that the deleted sequence in the promoter driving the trimethoprim resistance gene in pICOt had no significant influence on the fitness of the plasmid.

### Undigested pICOz, pICOt and pICOt2 run at a high molecular weight

We previously observed for pUCmu and pICOz that the undigested plasmids were running much higher than expected on a 1% Agarose/TAE gel [3]. To investigate if this is true also for pICOt and pICOt2, we digested pICOz, pICOt and pICOt2 with SpeI and ran the digested and undigested plasmids on a gel (Fig. 2).

**Fig. 2.**
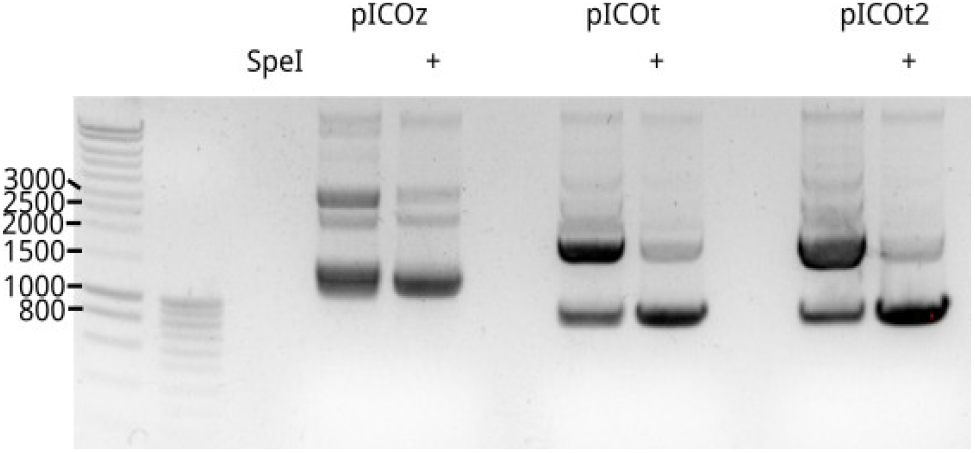
Comparison of undigested and SpeI digested pICOz, pICOt and pICOt2 plasmids on a 1% agarose/TAE gel. Left: Eurogentec SmartLadder and Eurogentec SmartLadder SF for size reference.

The higher-running undigested plasmids could be catenated – chains of interlocking plasmids. Catenated plasmids can be formed during plasmid replication [6], [7]. This seems to be something uniquely specific for our small plasmids (pUCmu (Ampicilin resistance; 1669 bp), pICOz (Zecoin resistance; 1185 bp) and pICOt2 (903 bp)). When something is cloned in these plasmid backbones, making them bigger, the normal coiled and supercoiled patterns appear [3]. Another possibility is that the very small plasmids simply do not coil normally and take more space than they should while running in the gel.

### Future perspectives

The 903 bp pICOt2 plasmid consists of a 237 bp selection marker (∼26%) and 616 bp minimal pUC origin of replication (ori; ∼68%), with the remaining ∼5% being the SpeI restriction enzyme site and the minimal promoter driving expression of the selection marker (49 bp). There is thus very little extra room for further miniaturization. There is a peptide (dcr1) which induces colistin resistance. This potential selectable marker is only 26 amino acids long (81 bp including stop codon). A colistin resistance pICOc plasmid could thus theoretically be as small as 747 bp. The pUC ori is already smaller than the original, based on a lucky accidental deletion in the generation of the pICOz plasmid [3]. “*There are no mistakes, just happy little accidents*” (Bob Ross) [8].

There are still smaller origins of replication, for example a 220 bp minimal essential origin of replication from pSC101 [9]. This origin of replication will however require a different helper plasmid or *E. coli* strain expressing the initiator protein RepA. An even smaller ori, also dependent on an external expression of Rep, has been engineered from the ColE2 ori. This ColE2-P9 minimal ori can be as small as 32-37 bp [10]. Plasmids with this minimal ori can for example be propagated in the so-called DIAL *E. coli* strains that express the Rep protein from their chromosomal DNA [11]. If that restriction would be acceptable, we could thus theoretically shrink a pICOt2-derived plasmid to approximately 319 bp, and a hypothetical pICOc-derived plasmid with this minimal ori would be only 163 bp. Potentially, the plasmid would have to be slightly bigger since we currently use a terminator sequence within the pUC ori to terminate the transcription of the selection marker. The smallest terminator sequence available at iGem is 34 bp (for example part BBa_B1002). Despite the theoretically impressive size advantages, these hypothetical plasmids with the minimal ori would however lose several of the advantages of the pICO series of plasmids, such as being high-copy and self-replicating in regular *E. coli* strains thanks to the pUC ori. The greatest opportunities for further size optimizations are thus in systematic deletions of parts of the minimal pUC ori and hope for more “happy accidents” by deleting other non-essential parts of the pUC ori.

## Materials and methods

### Synthetic generation of pICOt

The pICOt was designed *in silico* using UGene [12] from the pICOz sequence [3] by replacing the zeocin resistance gene with the 237 bp trimethoprim resistance gene dfrB2c (GenBank: ALZ46148.1) [13]. We also introduced several silent mutations in the resistance gene to eliminate restriction enzyme sites. The complete pICOt sequence was generated synthetically by IDT as a gBlock™. An additional opportunity to further reduce the plasmid size by 20 bp was to delete the sequence upstream of the -35 region in the AmpR promoter. The gBlock was consequently amplified with the primers pICOt-R : GAACACTAGTGTTTTTCCATAG and pICOt-F : GGTAACTAGTTATGTATCCGC for the complete gBlock or pICOt2-F : GGTAACTAGTAATAACCCTGATAAATGCTTC for the gBlock sequence with a minimal AmpR promoter. The PCRs were run with the high-fidelity Q5 DNA polymerase (New England Biolabs). The PCR products (pICOt and pICOt2) were subsequently cut with SpeI (Promega) and ligated with T4 DNA ligase (Promega) before being heat shock transformed into MC1061 *E. coli*.

### Evaluation of trimethoprim selection

Trimethoprim (92131-1G, Merck) was dissolved at a stock concentration of 50 mg/ml in DMSO and kept in a -20 °C freezer. *E. coli* MC1061 were transformed with the ligated pICOt and pICOt2 products and grown under 3 different conditions: regular (“Miller”) LB (10 g/L NaCl, 5 g/L yeast extract and 10 g/L tryptone) with 50 or 100 μg/ml trimethoprim, or low-salt (“Lennox”) LB (5 g/L NaCl, 5 g/L yeast extract and 10 g/L tryptone) with 50 μg/ml trimethoprim.

### Plasmid purification and validation

The plasmids were purified using a Macherey-Nagel plasmid miniprep kit, and Sanger sequenced using Eurofin (Mix2seq kit ON).

### Material availability

The plasmids pICOt (LMBP 13888) and pICOt2 (LMBP 13889) are available via BCCM/GeneCorner (https://www.genecorner.ugent.be/).

## Acknowledgments

We thank enthusiastic readers of our preprint publication for suggestions for further improvements and miniaturizations : Shyam Bhakta (Rice university, Houston, Texas) for suggesting the tiny 32 bp ColE2-derived ori, and Vuong Le (University of Copenhagen, Copenhagen, Denmark) for suggesting the tiny colistin resistance peptides.

## Notes

https://bccm.belspo.be/catalogues/lmbp-plasmids-plasmid-details?NM=pICOz

### Competing Interest Statement

The authors have declared no competing interest.

### Summary of Updates

We have adopted the manuscript based on feedback given by biorxiv preprint readers (figure, discussion/future perspectives). Independent researchers have already received the material generated here to "play" with and we expect this dynamic form of interaction with the wider community will shape the final form of this paper.

